# Doxorubicin induces caspase-mediated proteolysis of K_V_7.1

**DOI:** 10.1101/259242

**Authors:** Anne Strigli, Christian Raab, Sabine Hessler, Tobias Huth, Adam J. T. Schuldt, Christian Alzheimer, Thomas Friedrich, Paul W. Burridge, Mark Luedde, Michael Schwake

**Author notes:** Present address and correspondence should be addressed to: Michael Schwake Department of Neurology, Northwestern University Feinberg School of Medicine 303 East Chicago Avenue Chicago, IL 60611-4296. equal contribution (Anne Strigli, Christian Raab).

## Abstract

The voltage-gated potassium channel K_v_7.1 (KCNQ1) co-assembles with KCNE1 to generate the cardiac potassium current *I*_Ks_. Gain- and loss-of-function mutations in *KCNQ1* are associated with atrial fibrillation and long-QT (LQT) syndrome, respectively, highlighting the importance of modulating *I*_KS_ activity for proper cardiac function. On a post-translational level, *I*_KS_ can be regulated by phosphorylation, ubiquitination and sumoylation. Here, we report proteolysis of K_v_7.1 as a novel, irreversible posttranslational modification. The identification of two C-terminal fragments (CTF1 and CTF2) of K_v_7.1 led us to identify an aspartate critical for the generation of CTF2 and caspases as responsible for mediating K_v_7.1 proteolysis. Activating caspases by apoptotic stimuli significantly reduced K_v_7.1/KCNE1 currents, which was abrogated in cells expressing caspase-resistant K_v_7.1 D459A/KCNE1 channels. An increase in cleavage of K_v_7.1 could be detected in the case of LQT mutation G460S, which is located adjacent to the cleavage site. Application of apoptotic stimuli or doxorubicin-induced cardiotoxicity provoked caspase-mediated cleavage of endogenous K_v_7.1 in human cardiomyocytes. In summary, our findings establish caspases as novel regulatory components for modulating K_v_7.1 activity which may have important implications for the molecular mechanism of doxorubicin-induced cardiotoxicity.

**Non-standard Abbreviations and Acronyms:**

Cam: calmodulin
EBC: equilibrium buffer content
LQT syndrome: long QT syndrome
NRVM: Neonatal rat ventricular cardiomyocytes
hiPSC-CMs: human induced pluripotent stem cell-derived cardiomyocytes

## Introduction

Voltage-gated potassium channels (K_v_) form a protein class comprising forty members in humans, which can be grouped into twelve families [1]. Among these, the K_v_7 (KCNQ) family has attracted special attention since mutations in the five *KCNQ* genes cause heritable diseases, highlighting their physiological importance [2,3]. Dominant-negative mutations in the gene of K_v_7.1 are associated with cardiac arrhythmias contributing to LQT syndrome [4], whereas patients carrying loss-of-function mutations on both alleles additionally suffer from severe congenital hearing loss [5]. Gain-of-function mutations were found in patients with a form of autosomal dominant atrial fibrillation [6], highlighting the important functions of K_v_7.1 in the heart and inner ear. In both tissues, the α-subunit K_v_7.1 co-assembles with the β-subunit KCNE1, composing an ion channel conducting the slow component of the delayed rectifier potassium current, *I*_Ks_, which is indispensable for shaping the cardiac action potential [7,8]. Another important physiological function of K_v_7.1 was discovered by genome-wide association studies (GWAS), in which single nucleotide polymorphisms (SNP) in the *KCNQ1* locus were associated with type 2 diabetes in several populations [9,10]. In non-excitable (e.g., polarized thyroid, intestinal and tracheal epithelial) cells, K_v_7.1 associates with the β- subunits KCNE2 and KCNE3, respectively. The latter β-subunits, in contrast to KCNE1, reduce the voltage-dependent gating of the outwardly rectifying K_v_7.1 α-subunit, resulting in constitutively open channels. The K_v_7.1/KCNE2 channel complex has been described to be crucial for thyroid hormone biosynthesis [11], whereas K_v_7.1/KCNE3 heteromers play an important role in chloride secretion across tracheal and intestinal epithelia [12]. In the intestine, transposon-based forward mutagenesis genetic screens identified the *KCNQ1* gene as a cancer susceptibility gene [13] and low expression of K_v_7.1 was found in patients with colorectal cancer [14]. However, the role of K_v_7.1 in cancer development has yet to be established.

Channels formed from K_v_7 α-subunits share some structural features with *Shaker*-related K_v_ channels, such as a common core structure of six transmembrane domains (S1-S6) including a voltage-sensing domain (S1-S4) and a pore domain (S5-S6) [2]. One striking difference, however, is the presence of a large cytoplasmic C-terminal domain in K_v_7 channels, which is important for gating, assembly and intracellular trafficking of the channel and comprises four helical domains (A-D) (reviewed in Haitin et al. [15]). Whereas helices A and B mediate calmodulin binding [16], helices C and D form the subunit interaction (SI) domain [17], which consists of a bipartite coiled-coil motif that is crucial for subunit-specific interaction and tetramerization of the Kv7 α-subunits [18,19]. For K_v_7.1, it has also been shown that the A-kinase anchoring protein yotiao binds to helix D, leading to the recruitment of protein kinase A (PKA) and protein phosphatase 1, thereby forming a macromolecular complex important for regulating *I*_Ks_ activity [20]. In addition to PKA-mediated phosphorylation, ubiquitination and sumoylation have been reported to be important for regulation of K_v_7.1 activity at the posttranslational level [21,22].

Anthracyclines such as doxorubicin are well-stablished and effective anti-neoplastic agents, commonly used for the treatment of cancers. However, doxorubicin treatment has cardiotoxicity as a severe side-effect, which can lead to QT prolongation and heart failure [23,24]. Increased production of reactive oxygen species (ROS) and the induction of mitochondrial dysfunction are well described molecular mechanisms of doxorubicin-induced cardiomyopathy, resulting in an activation of apoptotic pathways, finally leading to caspase activation [25]. Dexrazoxane, a cardioprotective agent, acts in the same pathway by chelating Fe^2+^ ions more effectively than doxorubicin. While Fe^2+^ ions bound to doxorubicin are efficiently oxidized to Fe^3^+, resulting in release of electrons, which are rapidly transferred to produce ROS, the redox activity of dexrazoxane on Fe^2^+ ions is much lower, so that the Fe^2+^-scavenging activity and the consequently reduced ROS production by dexrazoxane have been suggested to underlie the alleviating effect on cardiotoxicity [26].

Caspases are cysteine proteases, which specifically cleave their substrates at the C-terminal side of an aspartic acid and play important roles in numerous aspects of physiology such as apoptosis, aging, development and inflammation [27]. Caspases are well known for their executive role in apoptosis and can be grouped into initiator (caspase 2, 8-10) and effector caspases (caspase 3, 6 and 7) [28]. Activation of caspases is triggered either by the extrinsic pathway, mediated by ligand binding to death receptors and activation of the initiator caspase-8, or by the intrinsic pathway [28]. In the latter case, mitochondrial membranes are permeablized by the pro-apoptotic proteins BCL-2 and BAX, leading to a loss of mitochondrial transmembrane potentials and to the release of other pro-apoptotic proteins such as cytochrome C into the cytosol, which results in the activation of the initiator caspase-9. Activated caspases 8 and 9 specifically cleave effector caspases, which finally execute apoptosis [28]. With the exception of caspase 14, all other caspases in humans have been implicated in inflammation [28]. Moreover, altered caspase expression levels have been correlated with ageing [29] and heart failure [30,31]. Recently, it has become evident that caspases also have non-apoptotic and non-inflammatory functions, such as regulation of long-term depression [32] or organelle removal during terminal differentiation [33].

In the present study, we identify K_v_7.1 as a novel substrate for caspases, which may have important implications for understanding the role of K_v_7.1 in cardiac arrhythmias and its function as a tumor suppressor. Our data suggest that caspase-mediated proteolysis of K_v_7.1 leads to decreased K_v_7.1-mediated currents, representing a novel regulatory mechanism for modulating K_v_7.1 channel activity. Furthermore, we show that K_v_7.1 cleavage is induced upon administration of doxorubicin, which efficiently activates caspase 3 in human cardiomyocytes. To our knowledge, K_v_7.1 is the first example of a voltage-gated potassium channel that acts as a substrate for caspases.

## Results

### Proteolysis of K_v_7.1 produces C-terminal fragments

We and others [34,35] have noted the occurrence of K_v_7.1 C-terminal fragments in transfected cells, when antibodies directed against epitopes on the K_v_7.1 C-terminus were used, which prompted us to further analyze the specificity of these fragments. We found C-terminal fragments of the full length K_v_7.1 channel in lysates derived from transiently transfected cells with human or murine K_v_7.1 cDNA constructs (Fig. 1A). Two fragments with a molecular mass of about ~40 kDa and ~28 kDa can be detected in addition to the full-length form of K_v_7.1 at ~70 kDa, when an antibody directed against a C-terminally derived peptide of K_v_7.1 was used (Fig. 1A and Fig. S1A). Next, we demonstrated the specificity of the K_v_7.1 antibody by the absence of signals in various tissues derived from K_v_7.1 deficient mice, which do not show detectable K_v_7.1 in immunoblots (Fig. S1B). As expected, we found the highest expression of Kv7.1 in murine heart, when compared to kidney and pancreas, further supporting the specificity of the antibody (Fig. S1B). C-terminal fragments of K_v_7.1 were also detectable in cells transfected with a cDNA construct of human K_v_7.1 tagged at the C-terminus with a MYC epitope (Fig. 1B). Immunoblots with either the MYC or the K_v_7.1 antibodies resulted in the same pattern. Next, we analyzed neonatal rat ventricular cardiomyocytes (NRVMs) for endogenous expression of K_v_7.1. Again, we found immunoreactive bands resembling full-length K_v_7.1 (~70 kDa) and bands at ~40 kDa and ~28 kDa, albeit with low intensity (Fig. S1C). Since the expression of an N-terminally tagged version of K_v_7.1 also revealed three complementary bands at ~70 kDa, ~44 kDa and ~30 kDa (data not shown), we concluded that K_v_7.1 is cleaved twice at its C-Terminus, resulting in three fragments: The N-terminal fragment comprising the cytoplasmic N-terminus and the membrane-embedded part of the protein and two C-terminal fragments showing immunoreactivity with the K_v_7.1 antibody, which we termed CTF1 (~40 kDa) and CTF2 (~28 kDa) (Fig. S1A).

**Figure 1:**
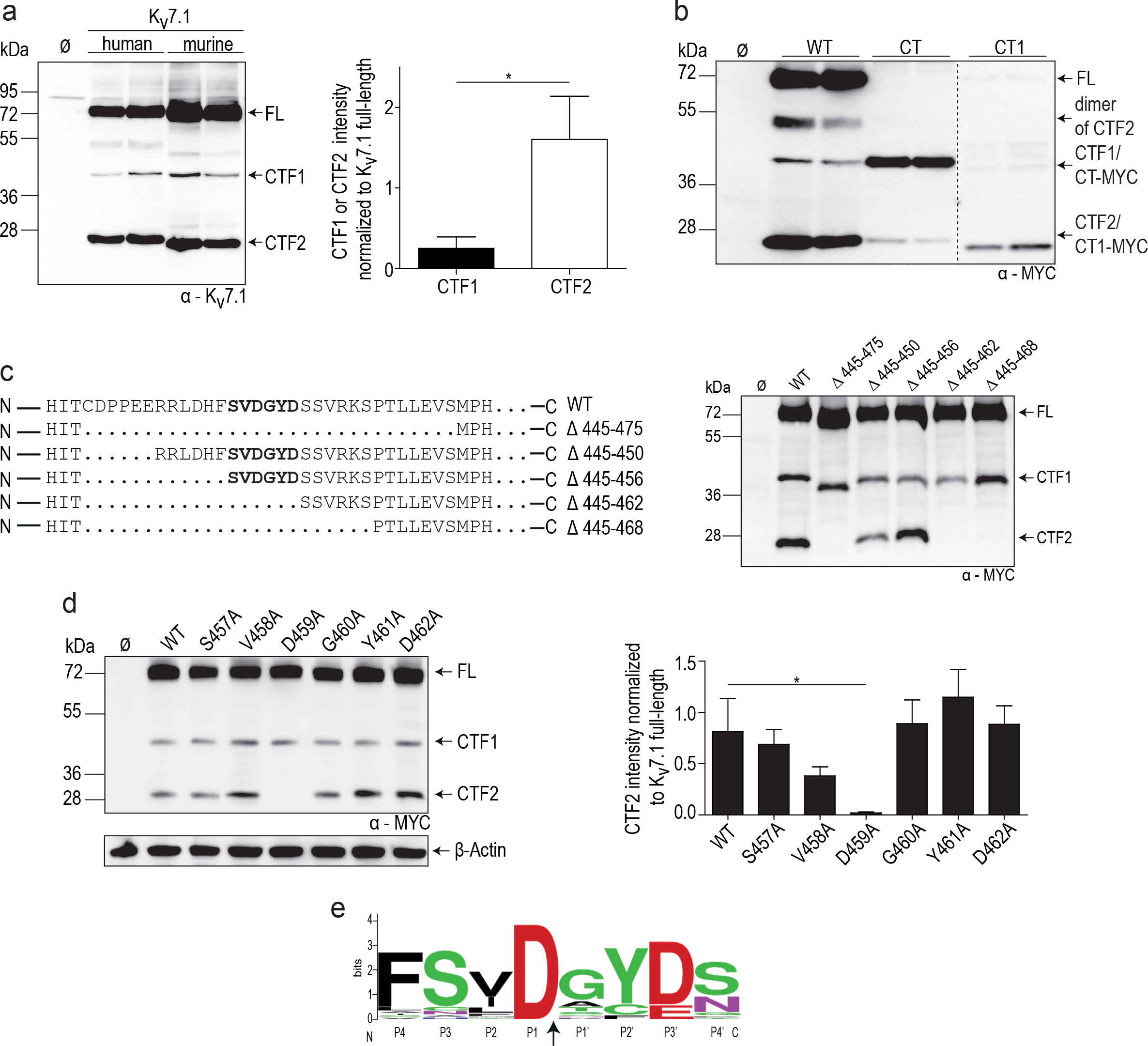
K_v_7.1 is cleaved at aspartate 459. **A**, Western blot analysis of HeLa cell lysates overexpressing human and murine K_v_7.1 constructs. Untransfected cells (Ø) served as negative control (left panel). Densitometric analysis of 4 independent experiments of the CTF1 or CTF2 band intensity normalized to the full-length K_v_7.1 band intensity. Statistics were tested with two-tailed Mann-Whitney U test. (right panel). **B**, Immunoblot analysis of lysates derived from HeLa cells overexpressing indicated constructs. Untransfected cells (Ø) served as negative control. **C**, Schematic illustration of different deletion constructs in the linker region between helix A and B. (left panel). HeLa cell lysates overexpressing indicated constructs analyzed by western blot. Untransfected cells (Ø) served as negative control (right panel). **D**, Alanine scan of position 457 to 462. Lysates from HeLa cells overexpressing indicated constructs were used for western blot analysis. Untransfected cells (Ø) served as negative control (left panel). Densitometric analysis of 6 independent experiments of CTF2 band intensity normalized to K_v_7.1 full length band intensity. Statistics were tested with Kruskal-Wallis test followed by Dunn’s multiple comparison test. (right panel). **E**, Cleavage site sequence logo of K_v_7.1 derived from 32 different species. Logo was created with weblogo.berkley.edu and adapted accordingly. The arrow indicates the cleavage site. (A) anti-K_v_7.1 antibody. (**B**-**D**) anti-MYC antibody.

To determine the cleavage sites, we generated two K_v_7.1 constructs: one comprising the complete C-terminus (CT, starting with glycine at position 348), and the other beginning at helix B (CT1 beginning with amino acid 505) (Fig. S1A). After expression of these fragments in HeLa cells, we compared the size of the resulting bands of CT and CT1 with the C-terminal fragments CTF1 and CTF2 (Fig. 1C). CT and CTF1 migrate at the same molecular weight, whereas CTF2 runs slightly slower than the CT1 construct. From these data, we conclude that the cleavage site for the generation of CTF1 resides between the end of the transmembrane domain S6 and helix A, while the cleavage site for CTF2 likely lies within the loop region between helices A and B in the C-terminus of K_v_7.1.

Since CTF2 is the most abundant C-terminal fragment (Fig. 1A, B, S1C), we focused our analysis on the region between helices A and B. We generated five K_v_7.1 constructs with varying deletions within this linker region (Fig. 1C). Immunoblot analysis of lysates derived from cells expressing these constructs revealed that a stretch of six amino acids (S^457^VDGYD^462^) was essential for the occurrence of CTF2 (Fig. 1C). Next, we performed an alanine scan over the SVDGYD region to define the precise cleavage site. Mutating the aspartate at position 459 to an alanine resulted in a complete loss of CTF2 generation, whereas mutating the other five residues (including Asp-462) appeared to have no significant effect (Fig. 1D). These data demonstrate that K_v_7.1 is cleaved within the SVDGYD motif at position D459 by a protease, which requires an aspartate. This aspartate is highly conserved within K_v_7.1 protein sequences derived from over twenty different species (Fig. 1E and Fig. S2). However, K_v_7.1 is the only member within the Kv7 family carrying an aspartate at this position (Fig. S1D).

### K_v_7.1 is cleaved by caspases upon apoptosis

To identify the protease responsible for the generation of CTF2, we searched the Merops Database for proteases that require an aspartate in the cleavage site [36]. Since caspases critically depend on an aspartate at the P1 position [27], we used two caspase inhibitors (Q-VD-OPH and Z-VAD(OMe)-FMK) to determine whether caspases are responsible for the generation of CTF2. Both compounds effectively inhibited the generation of CTF2 but not CTF1 (Fig. 2A and S3). To activate caspases, we induced apoptosis in cells overexpressing wild type K_v_7.1 and the D459A mutant by applying staurosporine, a non-selective protein kinase inhibitor widely used as a proapoptotic stimulus. Whereas wild type K_v_7.1 was efficiently proteolysed, the D459A mutant appeared resistant to staurosporine treatment (Fig. 2B), demonstrating that the D459A mutant is insensitive to staurosporine-induced caspase activation and cleavage. Next, we asked whether the generation of the CTF2 was dependent on the full-length K_v_7.1 α-subunit. We therefore expressed the CT construct (Fig. S1A) under control and apoptotic conditions. Again, an increase in CTF2 generation could be observed upon staurosporine treatment (Fig. 2C) indicating that cleavage can occur independent from the membrane-embedded part of the Kv7.1 protein. To gain deeper insight into the involvement of caspases in K_v_7.1 proteolysis, we applied staurosporine and the specific caspase 8 inhibitor II to cells expressing K_v_7.1. As shown in Fig. 3A, inhibition of caspase 8 efficiently blocked activation of downstream caspase 3 and the generation of CTF2 dose-dependently, demonstrating that the staurosporine-induced CTF2 generation is mediated by caspases.

**Figure 2:**
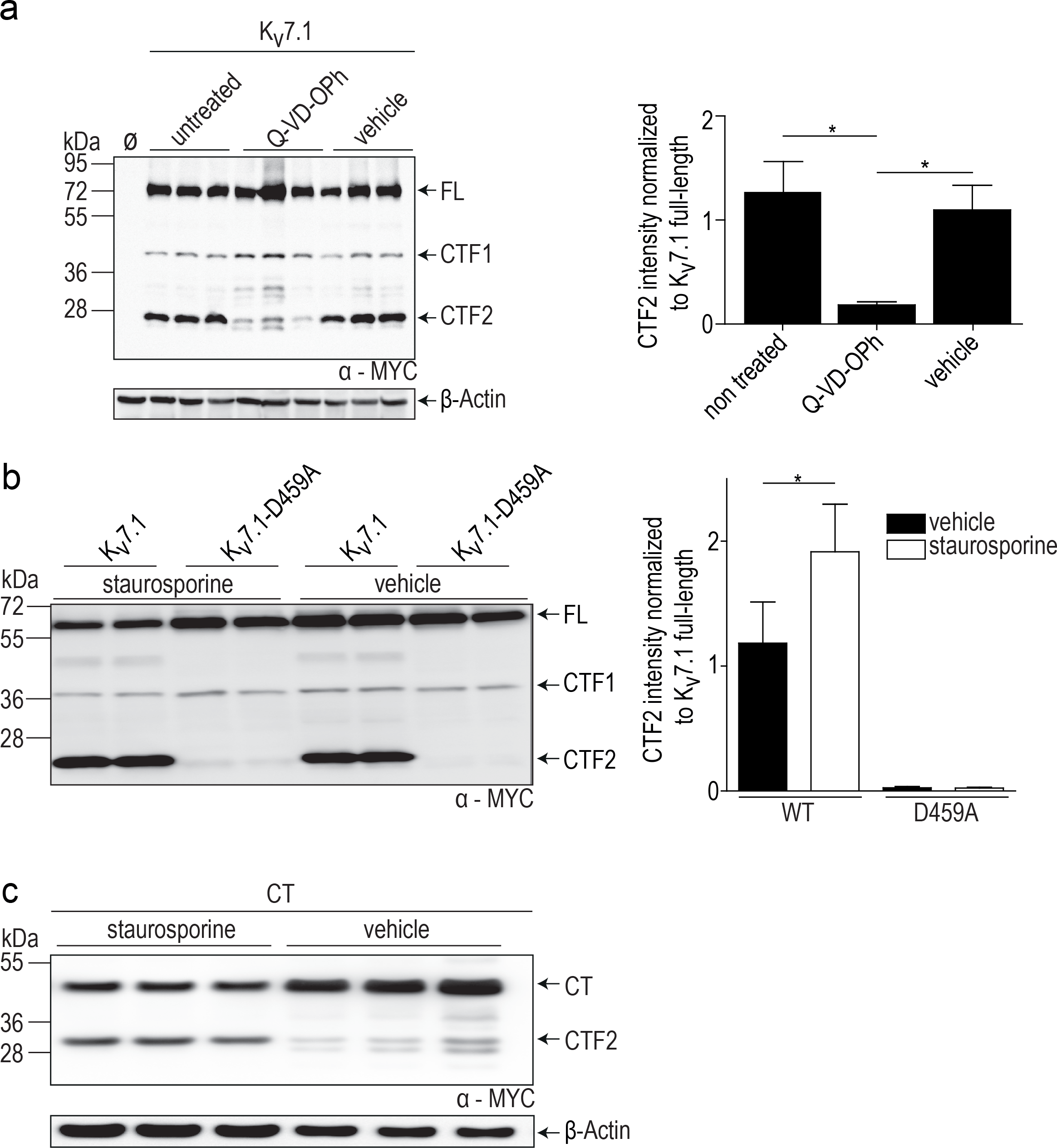
Cleavage of K_v_7.1 occurs during apoptosis. **A**, Western blot analysis of Cos7 cells overexpressing K_v_7.1-MYC treated for 12 hours with 450 nmol/L Q-VD-OPh. Untransfected (Ø), non-treated and vehicle-treated cells served as negative controls (left panel). Densitometric analysis of 5 independent experiments of CTF2 band intensity normalized to K_v_7.1 full length band intensity. Statistics were tested with Kruskal-Wallis test followed by Dunn’s multiple comparison test (right panel). **B**, Lysates derived from HeLa cells expressing K_v_7.1-MYC and K_v_7.1-D459A-MYC constructs treated with 1 μmol/L staurosporine for 8 hours analyzed by immunoblotting. Untransfected (Ø) and vehicle-treated cells served as negative controls (left panel). Densitometric analysis of 5 independent experiments of the CTF2 band intensity normalized to K_v_7.1 full-length band intensity. Statistics were tested with Kruskal-Wallis test followed by Dunn’s multiple comparison test (right panel). **C**, Western blot analysis of HeLa cell lysates overexpressing CT-MYC construct treated for 8 hours with 1 μmol/L staurosporine. Vehicle-treated cells served as negative controls. (**A**-**C**) anti-MYC antibody, (**A,C**) anti-β-actin antibody.

**Figure 3:**
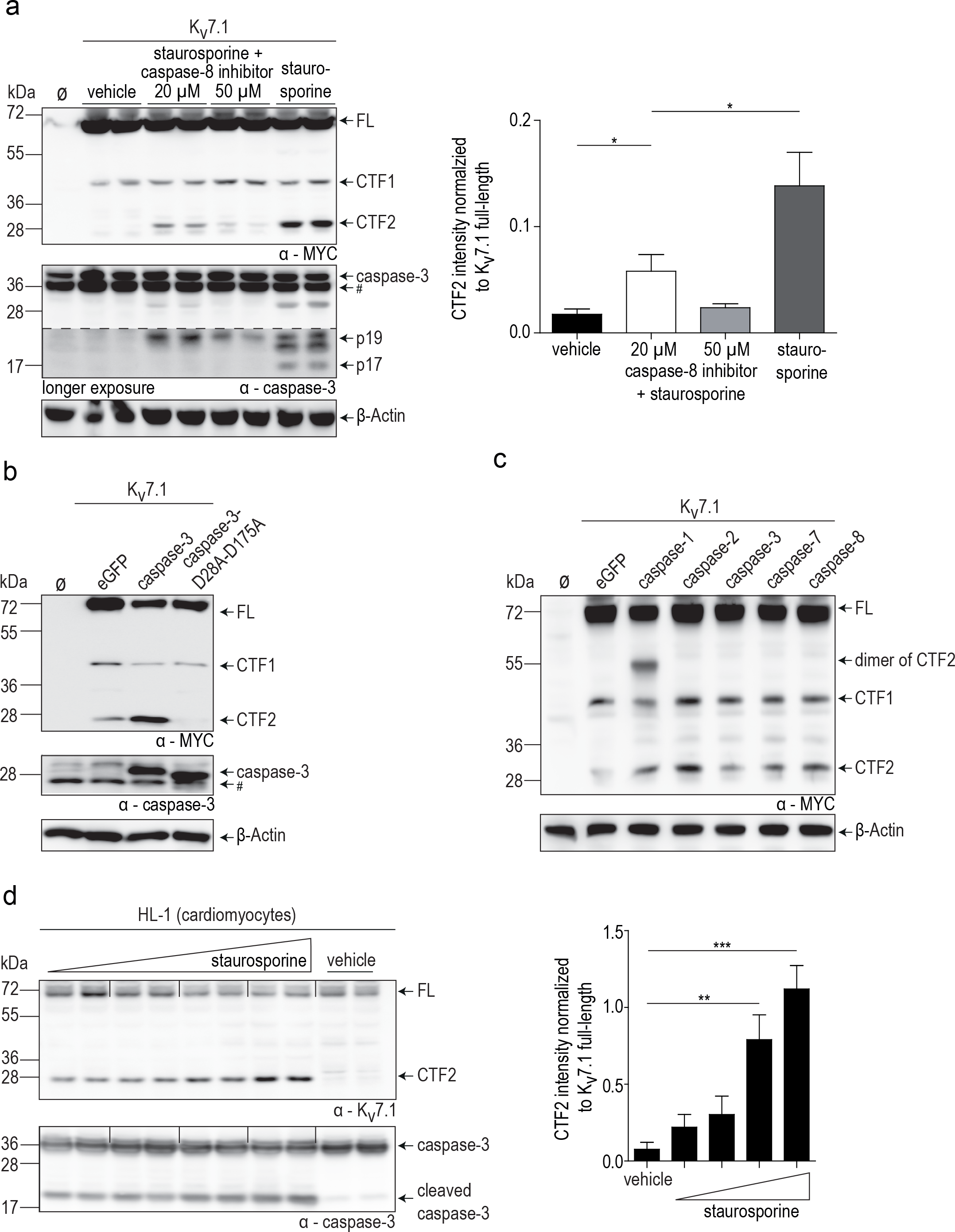
Caspases are responsible for the generation of CTF2. **A,** Western Blot analysis of lysates derived from HEK 293T cells stably expressing K_v_7.1-MYC treated with either 1 μmol/L staurosporine for 6 hours or treated with 1 μmol/L staurosporine for 6 hours and pre-treated for 2 hours with 20 or 50 μmol/L of a caspase-8 inhibitor II. Untransfected (Ø) and vehicle-treated cells served as negative controls (left panel). Densitometric analysis of 3 independent experiments of the CTF2 band intensity normalized to K_v_7.1 full length band intensity. Statistics were tested with Kruskal-Wallis test followed by Dunn’s multiple comparison test. (right panel). # indicates nonspecific binding of the antibody. **B,** MCF-7 lysates expressing K_v_7.1-MYC and caspase-3 or caspase-3-D28A-D175A analyzed by immunoblot. Untransfected (Ø) and eGFP-transfected cells served as negative controls. # indicates nonspecific binding of the antibody. **C,** Lysates of HEK 293T cells stably expressing K_v_7.1 and co-expressing indicated caspases analyzed by immunoblot. Untransfected (Ø) and eGFP transfected cells served as negative controls. **D,** Lysates of HL-1 cells treated for 6.5 hours with 0.5, 1, 1.5 and 2 μmol/L of staurosporine analyzed by immunoblot. Vehicle-treated cell lysates served as negative control (left panel). Densitometric analysis of CTF2 band intensity normalized to K_v_7.1 full-length band intensity of 5 independent experiments. Statistic were tested with One-Way-ANOVA followed by Bonferroni’s Multiple Comparison test. (right panel). **(A – C)** anti-MYC antibody. anti-β-actin antibody. **(a,b,d)** anti-caspase-3 antibody. **(D)** anti-K_v_7.1 antibody.

To address the question whether caspase 3, one of the major effector caspases, is solely responsible for K_v_7.1 cleavage at position D459, we used a human breast carcinoma MCF-7 cell line, which is deficient for caspase 3 [37]. Overexpression of K_v_7.1 in MCF-7 cells still resulted in the generation of CTF2 albeit to a lower extent (Fig. 3B, eGFP labeled lane). Whereas co-transfection of K_v_7.1 together with wild-type caspase-3 restored CTF2 production, the overexpression of an inactive form of caspase-3 led to a reduction of CTF2 levels (Fig. 3B) likely by protecting K_v_7.1 from endogenous caspases. To determine whether all caspases cleave K_v_7.1 to the same extent, we co-expressed K_v_7.1 with at least one caspase of each group, namely caspase 1, 2, 3, 7 and 8. Immunoblots revealed that all tested caspases are able to generate CTF2 (Fig. 3C). Stronger cleavage could be observed by overexpression of the initiator caspase-1, -2 and -8, which might be due to an activation of downstream effector caspases. Nevertheless, these data suggest that K_v_7.1 is a substrate of all analyzed caspases.

To demonstrate that endogenous K_v_7.1 undergoes caspase-mediated proteolysis, we treated murine cardiac muscle cells (HL-1 cells [38]) with increasing concentrations of staurosporine, confirming a dose-dependent occurrence of CTF2 (Fig. 3D). Notably, full length K_v_7.1 channel was efficiently cleaved at higher staurosporine concentrations. In summary, our data strongly suggest that K_v_7.1 is cleaved by caspases at an aspartate at position 459.

### Functional impact of caspase-mediated proteolysis on wild-type K_v_7.1 and identification of LQT1 mutations modulating K_v_7.1 proteolysis

To analyze the functional impact of caspase-mediated cleavage of K_v_7.1 upon induction of apoptosis, we co-transfected HEK 293T cells with wild-type K_v_7.1 and the D459A mutant together with KCNE1 and measured whole-cell currents under staurosporine treatment and control conditions (Fig. 4A). Drug treatment produced a significant reduction of K_v_7.1/KCNE1 currents, whereas currents generated by K_v_7.1 D459A/KCNE1 channels remained unaffected (Fig. 4B). Cells were harvested after patch-clamp measurements and were subjected to immunoblot analysis to probe for CTF2. Again, we were able to detect CTF2 in cells expressing wild type K_v_7.1 but not the D459A mutant, as shown in Fig. 4C.

**Figure 4:**
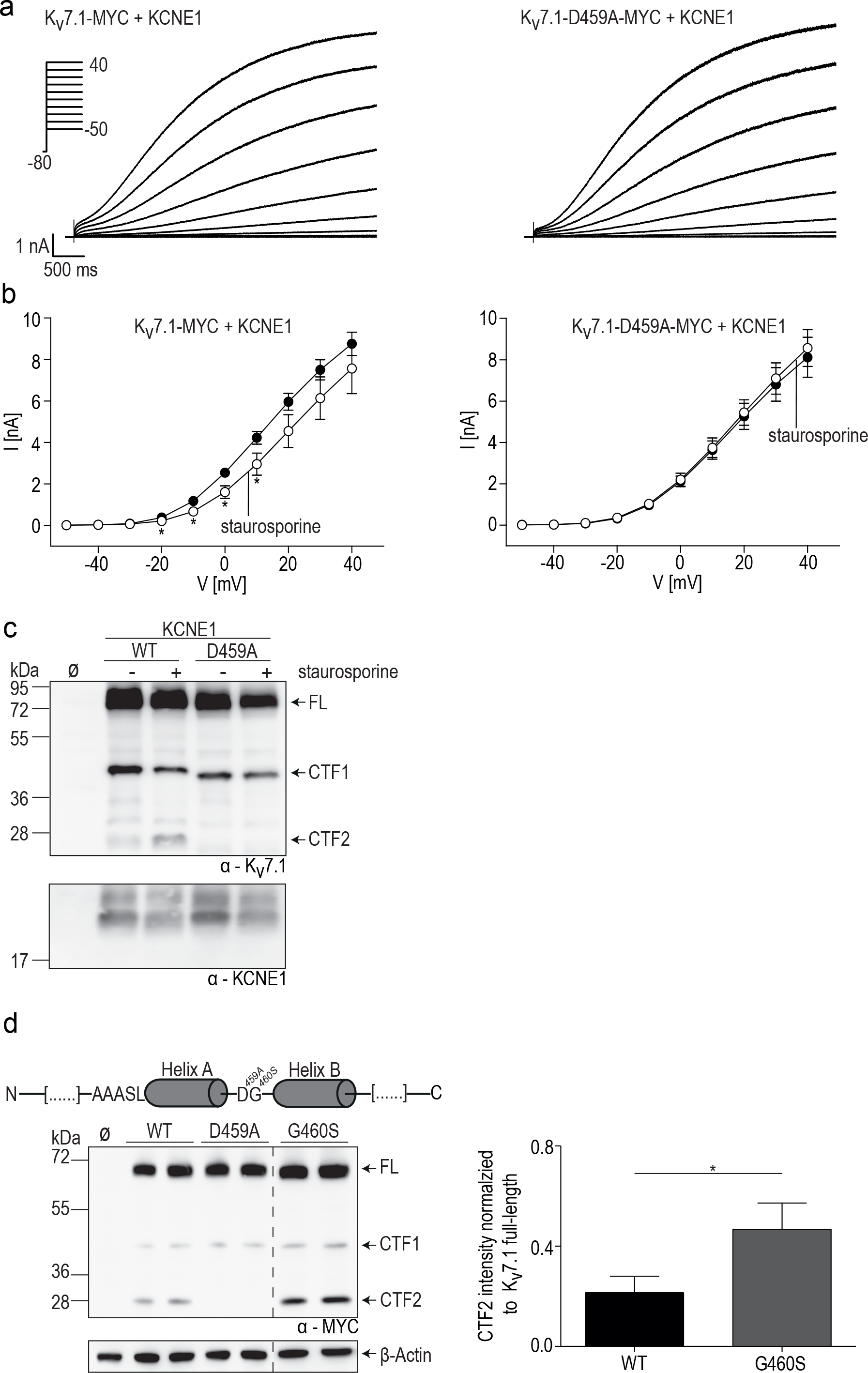
Cleavage of K_v_7.1 in physiology and pathophysiology. **A**, Representative current traces for K_v_7.1-MYC and K_v_7.1-D459A-MYC, both co-expressed with KCNE1. **B**, Mean currents amplitude was plotted versus voltage to obtain current-voltage (I-V) relationships in cells expressing K_v_7.1-MYC or K_v_7.1-D459A-MYC and KCNE1 treated with 500 nmol/L staurosporine for 10 – 12 hours. Statistics were tested with Two-Way ANOVA followed by Bonferroni posttests. **C**, Western blot analysis of cells used for patch-clamp recordings. **D**, Schematic illustration to highlight the position of G460 and calmodulin binding site in helix A (upper left panel). Western blot analysis of HeLa cell lysates overexpressing indicated constructs. Untransfected (Ø) and K_v_7.1-D459A transfected cells served as negative controls (lower left panel). Densitometric analysis of four independent experiments of CTF2 band intensity normalized to K_v_7.1 full-length band intensity. Statistics were tested with One-Way-ANOVA followed by Bonferroni’s Multiple Comparison test. (right panel). **(C)** anti-K_v_7.1 antibody, anti-KCNE1 antibody. **(D)** anti-MYC antibody, anti-β-actin antibody.

Next, we tested if disease-causing mutations in K_v_7.1 can interfere with the generation of CTF2 and focused on the LQT1 mutation G460S, which is located just one amino acid downstream of the aspartate residue important for K_v_7.1 proteolysis (Fig. 4D). Immunoblot analysis of the G460S mutant protein demonstrated significantly increased CTF2 levels when compared to wild type K_v_7.1, indicating that this LQT1 mutation renders K_v_7.1 more susceptible to caspase-mediated cleavage even under non-staurosporine treatment conditions (Fig. 4D). This observation is in line with the finding that this mutation causes a decrease in *I_Ks_*-like currents [39]. Other LQT1 mutations (D446E, P448L and R451W) in the proximity of the D459 residue did not affect CTF2 generation (data not shown).

### Doxorubicin induces K_v_7.1 proteolysis in stem cell-derived cardiomyocytes

It is well known that cancer treatment by the common antineoplastic doxorubicin is hindered by severe cardiotoxic side effects, and there is strong evidence in the literature that doxorubicin leads to caspase activation in cardiomyocytes [40]. In order to determine, whether interference with cardiac function by doxorubicin also involves caspase-mediated cleavage of Kv7.1, we treated human induced pluripotent stem cell-derived cardiomyocytes (hiPSC-CMs) with staurosporine and doxorubicin. Both compounds activated caspase 3 as indicated by the occurrence of cleaved-active forms and a reduction of the inactive zymogen of the protease (Fig. 5). CTF2 was detectable after compound treatments, but was absent from mock treated cells. However, doxorubicin treatment appeared to be more efficient in producing active caspase 3, which correlated with a higher abundance of CTF2 and a strong reduction of monomeric and tetrameric forms of Kv7.1 (Fig. 5). Interestingly, we detected a potentially dimeric form of Kv7.1 at 150 kDa in staurosporine and doxorubicin treated cells, suggesting that the caspase-mediated destruction of tetramers results in Kv.7.1 dimers in hiPSC-CMs (Fig. 5).

**Figure 5:**
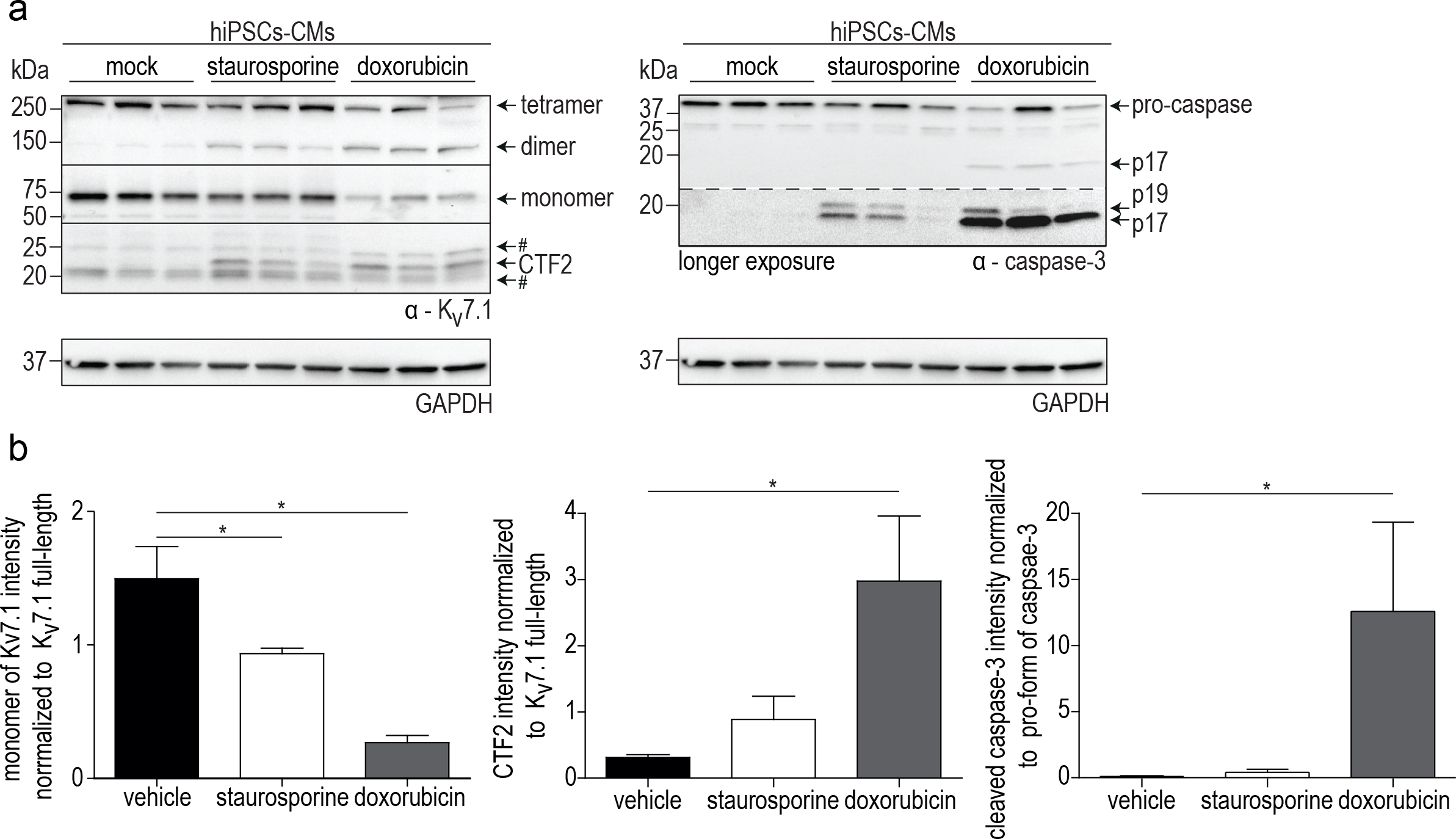
Doxorubicin induces cleavage of K_v_7.1 to CTF2. **A**, Immunoblot analysis of hiPSC-CMs treated either with vehicle, staurosporine (2 μM) for 4 hours or doxorubicin (10 μM) overnight. # indicates nonspecific binding of the K_v_7.1 antibody. **B,** Densitometric analysis of 3 independent experiments of band intensities of Kv7.1 monomer normalized to GAPDH (left panel), CTF2 to K_v_7.1 monomer (middle panel) and cleaved caspase 3 (p19 + p17) to the pro-form of caspase 3 (right panel). Statistics were tested with one-tailed Mann-Whitney U. **(A)** anti-K_v_7.1 antibody, anti-caspase-3 antibody and anti-GAPDH antibody were used.

## Discussion

To date, over 1500 caspase cleavage sites and substrates have been identified [41]. The vast majority of caspase-mediated cleavage occurs during apoptosis, but caspase function has also been demonstrated in non-apoptotic cellular responses [42], suggesting that caspases cleave a specific subset of substrates without inducing apoptosis. Here, we report K_v_7.1 as a novel substrate for caspases, representing to our knowledge the first example of a voltage-gated cation channel undergoing caspase-mediated proteolysis.

Cleavage of human K_v_7.1 by caspases occurs after an aspartate at position 459, which is located within the intervening loop between helices A and B in the channel`s large cytoplasmic C-terminal domain that serves as a scaffold for numerous protein-protein interactions involved in cellular signaling cascades [15]. Both helices appear to form a two-helical bundle, which is embraced by a CaM molecule as revealed by X-ray crystallography of recombinantly expressed CaM and the proximal C-terminus of K_v_7.1 [34]. In this study, the intervening loop, in which the caspase-mediated proteolysis of K_v_7.1 occurs, was deleted. Therefore, structural information about the cleavage site is missing.

Our functional analysis using patch-clamp strongly suggests that caspase cleavage entails loss-of-function of K_v_7.1. This finding is supported by numerous reports showing that the intracellular C-terminal domain of K_v_7.1 is responsible for channel tetramerization, trafficking and modulating the biophysical properties of the channel [15]. In line with this, many pathogenic LQT1 mutations have been mapped to the C-terminus of K_v_7.1 emphasizing the functional importance of this particular channel region [43]. Our finding that the LQT1 mutation G460S, which is adjacent to the aspartate 459 residue, is even more susceptible to caspase cleavage suggest a novel pathophysiologic mechanism for LQT1 mutations located in the C-terminus of K_v_7.1.

Based on the analysis of a number of cleavage sites, a general consensus motif of DXED-A/G/S/T has been proposed for apoptosis executioner caspases such as caspase 3 and 7, whereas caspases 2, 8, 9, and 10 and caspases 1, 4, 5 6, 14, prefer isoleucine/ leucine or tryptophan/tyrosine/valine instead of an aspartate at the first position, respectively [41]. Due to the overlapping specificity of caspases and the significantly different K_v_7.1 cleavage site (amino acid sequence: FSVD-G), it is difficult to predict which individual caspase cleaves K_v_7.1. Our co-expression data suggest that K_v_7.1 can be cleaved by all tested caspases including caspase 1, 2, 3, 7 and 8. In this group, caspase 1 was more efficient in K_v_7.1 processing, which is in agreement with the larger similarity of the preferred cleavage site [41].

Caspases have well-established functions in the execution of apoptosis as well as inflammation [28]. However, transiently active caspases have also been detected in non-apoptotic cells. For example, in neurons, caspases 3 and 9 are critical for long term depression (LTD) and AMPA receptor internalization [32]. In the heart, increased expression of caspase 1 was found in murine heart failure models and in patients with end-stage heart failure [31]. Analysis of mice with heart-targeted overexpression of caspase 1 and 3 further supported the notion that caspases contribute to heart diseases, likely based on an overlap of apoptotic and non-apoptotic functions [30,31]. Our finding that I_Ks_ is sensitive to caspase-mediated cleavage uncovers a novel molecular mechanism that may contribute to cardiac arrhythmias, which is strongly supported by an increased susceptibility of the LQT1 G460S mutant to proteolytic processing by caspases. Thus, it is likely that the reported 40% smaller current density of the G460S mutant, when compared to wild type I_Ks_, is at least partially due to an increased cleavage of the mutant [39]. Although several C-terminally located LQT1 mutations have been identified, which interfere with channel function by modulating CaM [16,44] or PIP_2_ [45] binding and/or disrupt assembly of functional I_Ks_ [15], for most of the C-terminal LQT mutations, the pathophysiological mechanism that leads to disease is unknown. Our data strongly suggest that susceptibility to caspase-mediated degradation should also be considered when analyzing these mutations.

Furthermore, our results demonstrate that doxorubicin treatment of hiPSC-derived cardiomyocytes efficiently induced caspase-mediated cleavage of K_v_7.1, suggesting that this pathway contributes to doxorubicin-induced cardiotoxicity. Interestingly, doxorubicin has also been shown to induce electrocardiogram abnormalities such as QT interval prolongations, which are often observed within the first day after chemotherapy [25]. These results have been confirmed in animal studies, showing that doxorubicin prolongs the cardiac action potential duration by specifically inactivating I_Ks_ but not I_Kr_, which both compose the delayed rectifier potassium current I_K_ [23]. In cardiomyocytes, I_Ks_ is mediated by a macromolecular complex formed by assembly of the pore-forming subunits K_v_7.1 with KCNE1 β-subunits, which are linked to the scaffolding protein yotiao/A-kinase anchoring protein 9 (AKAP-9) [20]. Yotiao binds to the distal part of the C-Terminus of K_v_7.1 and recruits PKA, protein phosphatase 1 (PP1), adenylate cyclase 9 (AC9) and phosphodiesterase PDE4D3 to the complex, allowing the control of the phosphorylation state of K_v_7.1, which is the molecular basis for the β-adrenergic regulation of I_Ks_ [20,46]. *In silicio* sequence analyses of yotiao predict several potential caspase cleavage sites, suggesting that this scaffold protein is also processed by caspases [41]. Thus, it is conceivable that caspase-mediated processing of K_v_7.1 and likely yotiao contributes to the prolongation of the QT interval mediated by doxorubicin. It will be important to determine the pathway, by which doxorubicin treatment leads to elevated caspase activity and K_v_7.1 cleavage. While it is widely accepted that doxorubicin-induced cardiotoxicity is due to the induction of mitochondrial dysfunction, resulting in an increased production of reactive oxygen species (ROS) in the cytoplasm and consequent activation of extrinsic and intrinsic apoptotic pathways, a more direct effect on ROS-mediated signaling by the oxidizing activity of doxorubicin on Fe^2+^ ions, as suggested by the alleviating effect of dexrazoxane, also needs to be considered. This will also clarify the issue of whether regulated C-terminal cleavage of K_v_7.1 is more generally involved in ROS-mediated cardiac responses.

In summary, the present study demonstrates caspase-mediated proteolysis of K_v_7.1. Posttranslational modifications such as phosphorylation, ubiquitination, sumoylation, palmitoylation and glycosylation have been reported for potassium channels [47]. According to our data, proteolysis of K_v_7.1 mediated by caspases is another important mechanism of post-translational modification of K_v_7.1, and we hypothesize that analysis of this regulation will forward understanding of the molecular mechanism of doxorubicin-induced cardiotoxicity.

## Methods

### Plasmids

Human K_v_7.1 and caspase cDNAs were subcloned in the expression vectors pFrog and pcDNA4/TO (Invitrogen, Waltham, USA), respectively. Mutations, deletions and tags for antibodies were constructed/introduced by recombinant PCR and verified by sequencing.

### Antibodies

The following antibodies were used: rabbit anti-β-actin (A2066, Sigma-Aldrich, St. Louis, USA), rabbit anti-caspase 3 (8G10, Cell Signaling, Cambridge, UK), rabbit anti-calmodulin (ab45689, Abcam, Cambridge, UK), mouse anti-GAPDH (MAB374, Millipore, Billerica, USA), rabbit anti-KCNE1 (APC-163, Alomone Labs, Jerusalem, Israel), rabbit anti-K_v_7.1 (ab77701, Abcam, Cambridge, UK), mouse anti-MYC (9B11, Cell Signaling, Cambridge, UK), goat anti-MYC (GTX29106, GeneTex Inc., Irvine, USA). The C-terminal anti-K_v_7.1 antibody was directed against the following peptide sequence TVPRRGPDEGS.

### Cell culture, transfection and inhibitors

Cell lines were grown in Dulbecco’s modified Eagle’s medium (DMEM, Thermo Fisher Scientific) supplemented with 10% fetal bovine serum (FBS, Biochrome, Berlin, Germany), 100 U/mL penicillin and 100 μg/mL streptomycin (both Carl Roth, Karlsruhe, Germany). For HL-1 cells, Claycomb medium (Sigma-Aldrich), supplemented with 10% FBS (Biochrome), 100 U/mL penicillin, 100 μg/mL streptomycin, 0.1 mM norepinephrine (Carl-Roth) and 2 mM L-Glutamine (Carl Roth) was used. All cells were kept at 37 °C and 5% CO_2_. Transient transfections were performed using TurboFect (Thermo Fischer Scientific) according to the manufacturer’s instructions, and stable cell lines were established using G418 for selection. Inhibitors were used as follows: Z-VAD-FMK (12 h, 100 μM, Promega, Fitchberg, USA), Q-VD-OPh (12 h, 450 nM, Merck Millipore, Billerica, USA), staurosporine (1 to 8 hours, 0.5 – 2 μM, Sigma-Aldrich), caspase-8 inhibitor II (2 h pretreatment, 20 or 50 μM, Calbiochem, Billerica, USA).

### Protein extraction, immunoprecipitation and immunoblot analysis

Cells were washed twice with phosphate buffered saline (PBS) and harvested in PBS, containing a protease inhibitor cocktail (Complete, Roche, Basel, Switzerland). After centrifugation, cell pellets were lysed in PBS/Complete (1% Triton X-100, Carl Roth) by sonication. After 1 h incubation on ice, samples were centrifuged, and supernatants were analyzed by SDS-PAGE. For co-immunoprecipitations, cells were lysed in EBC buffer/Complete (120 mM, NaCl 50 mM, Tris-HCl, 0.5% NP-40, pH 7.4 (all from Carl-Roth)) and treated as described above. Lysates were incubated with mouse □-MYC antibody at 4°C overnight. For precipitation, Protein G agarose beads were used. After thorough washing of the beads, protein complexes were released by denaturation. Samples were subjected to SDS-PAGE and transferred onto nitrocellulose membranes by tank blotting. Membranes were blocked and incubated overnight at 4°C in primary antibody solution followed by an incubation with the appropriate secondary antibodies conjugated to horseradish peroxidase (HRP). After thorough washing, bound antibodies were detected by chemiluminescence using a luminescent imager (LAS-4000, Fujifilm, GE Healthcare, Little Chalfont, UK). For quantifications, ImageJ software was used.

### Isolation of neonatal rat ventricular cardiomyocytes (NRVM)

Hearts of 1 - 2 days old Wistar rats were harvested and minced in buffer (120 mMol NaCl, 20 mMol HEPES, 8 mMol NaH_2_PO_4_, 6 mMol glucose, 5 mMol KCl, 0.8 mMol MgSO_4_, pH = 7.4). Subsequently, up to six digestion steps were carried out with 0.6 mg/ml pancreatin (Sigma-Aldrich) and 0.5 mg/mL collagenase type II (Worthington, Lakewood, USA) in sterile ADS buffer. Cardiomyocytes were purified from contaminating fibroblasts using a Percoll gradient centrifugation step. Finally, NRVMs were resuspended and cultured in DMEM containing 10% FBS, 100 U/mL penicillin, 100 μg/mL streptomycin and 1 % L-glutamine. Protein extractions were performed as described above.

### Human induced pluripotent stem cell culture and cardiac differentiation

Human iPSCs were routinely maintained in E8 medium supplemented with 10 μM Rho kinase inhibitor (Y27632; Biorbyt, Cambridge, UK) for the first 24 h after passage on 1:400 reduced growth factor Matrigel (Corning, Corning, USA). Cells were passaged ~1:15 every 3-4 days using 0.5 mM EDTA in Dulbecco’s PBS (DPBS; Corning, Corning, USA) after achieving ~80% confluence. Cell lines were used between passages 20 and 85. All cultures were routinely tested for mycoplasma using a MycoAlert PLUS Kit (Lonza, Basel, Switzerland). Cardiac differentiation was performed as described in [48]. Briefly, to initiate differentiation, medium was changed to CDM3, consisting of RPMI 1640 (Corning), 500 μg ml^−1^ *Oryza* sativa–derived recombinant human albumin (Oryzogen, Wuhan, China), and 213 μg ml^−1^ L–ascorbic acid 2–phosphate (Wako, Tokyo, Japan). For days 0–1, media was supplemented with 3 μM of the glycogen synthase kinase 3–β inhibitor CHIR99021 (Biorbyt, San Francisco, USA)^21,22^ and 10 ng ml^−1^ BMP4 (Peprotech, Hamburg, Germany). On day 1, media was changed to CDM3 and on d2 media was changed to CDM3 supplemented with 2 μM of the Wnt inhibitor Wnt–C59 (Biorbyt). Media was changed on day 4 and every other day thereafter with CDM3. Contracting cells were noted from day 7. At days 25-29, contracting cardiomyocytes were dissociated by incubating 20 minutes in DPBS followed by 6 minutes in TrypLE (Thermo Fisher Scientific), and then 60 minutes with 0.5 U/ml Liberase TH (Roche) in CDM3 media. Cells were replated in 40% FBS in CDM3 by combining 2-3 wells into one well for optimal viability of the cultures. After 2 days, media was changed back to CDM3 and exchanged every 2 days until analysis. and in supplementary information.

### Electrophysiology

Transfected cells were identified by using co-transfection of eGFP and imaging on an inverted fluorescence microscope (Axiovert 40, Zeiss, Jena, Germany) with a fiber optic-coupled light source (UVICO, Rapp OptoElectronic, Hamburg, Germany). Current signals were recorded in whole-cell patch-clamp mode at room temperature (22±1°C) two days after transfection. Recordings were started 3 min after whole-cell access was obtained. Data were sampled at 20 kHz and filtered at 5 kHz, using an Axopatch 200B amplifier in combination with a Digidata 1322A interface and pClamp10 software (all from Molecular Devices/MDS Analytical Technologies, Sunnyvale, USA). Electrodes were made from borosilicate glass (Harvard Apparatus, Edenbridge, UK or BioMedical Instruments, Zoellnitz, Germany), using a DMZ-Universal Puller (Zeitz, Munich, Germany). Pipette resistance in bath solution was 2.0-3.5 MΩ and access resistance was typically < 5 MΩ before series resistance compensation (75%). External solution contained (in mM): 145 NaCl, 4 KCl, 2 CaCl_2_, 2 MgCl_2_, 10 D-Glucose, 10 HEPES, adjusted to pH 7.4 with NaOH. The internal solution was composed of (in mM) 135 K-gluconate, 4 NaCl, 10 KCl, 5 Hepes, 5 EGTA, 2 Na_2_-ATP, 0.3 Na_3_-GTP (pH 7.25 with KOH). Cells were incubated with staurosporine at a concentration of 500 nM 10-12 hours before recordings. Chemicals were purchased from Sigma-Aldrich.

### Statistics

Data are shown as means ± SEM of n observations, as indicated. Statistical analyses were performed using unpaired one- or two-tailed student’s t-Test, Mann-Whitney-U, One-Way ANOVA and Bonferroni’s Multiple Comparison post hoc test, Kruskal-Wallis test followed by Dunn’s Multiple Comparison Test or Two-Way ANOVA followed by Bonferroni posttests, where applicable, using the GraphPad Prism 5 software (GraphPad, San Diego, USA). Normal distribution was tested with KS normality test, D’Agostino & Person omnibus normality test and Shapiro-Wilk normality test. Mann-Whitney-U and Kruskal-Wallis test were only applied if all three tests failed. p ≤ 0.05 was termed significant. * 0.01 ≤ p ≤ 0.05. ** 001 ≤ p < 0.01. *** p < 0.001.

## Acknowledgements

We thank Maike Langer, Vanessa Mangels and Marvin Murowski for excellent technical assistance; William C. Claycomb for the HL-1 cells and Jakob Völkl, Florian Lang and Karl E. Pfeifer for K_v_7.1-deficient murine tissues. This work was supported by the Deutsche Forschungsgemeinschaft (Grant SFB 877, B8 and Heisenberg Fellowship to M.S.).

